# The Olfm4-defined human neutrophil subsets differ in proteomic profile in septic shock

**DOI:** 10.1101/2022.03.08.483264

**Authors:** Hans Lundquist, Henrik Andersson, Michelle S. Chew, Jyotirmoy Das, Maria V. Turkina, Amanda Welin

**Author notes:** Corresponding author: Amanda Welin, Division of Inflammation and Infection, Department of Biomedical and Clinical Sciences, Lab1 Floor 12, Linköping University, SE-58185 Linköping, Sweden.

## Abstract

The specific granule glycoprotein olfactomedin-4 (Olfm4) marks a constitutive subset of neutrophils in humans, where 1-70% of peripheral blood neutrophils produce Olfm4. The proportion of Olfm4-high (Olfm4-H) neutrophils correlates with the severity of paediatric septic shock and could predict mortality in adult septic shock in previous studies, but it is not known whether and how the Olfm4-H neutrophils contribute to sepsis pathogenesis. The aim of this study was to decipher proteomic differences between the Olfm4-H and Olfm4-low (Olfm4-L) human neutrophil subsets at baseline and in the context of septic shock, hypothesizing that Olfm4 marks a neutrophil subset with a distinct proteomic profile, predisposing it for detrimental processes in sepsis. A novel protocol for the preparation of fixed, antibody-stained and sorted neutrophils for LC-MS/MS analysis of proteome was developed. In neutrophil subsets from healthy blood donors, 47 proteins had significantly higher abundance in the Olfm4-H population, and 62 proteins in the Olfm4-L population. Pathway enrichment analysis showed that the differences concerned proteins related to neutrophil degranulation, with *e.g.* Rab3d and a subunit of the vacuolar ATPase proton pump being more abundant in the Olfm4-H neutrophils, and the alarmin S100-A7, the major neutrophil chemotactic receptor CXCR1 and the antimicrobial peptide defensin alpha-4 being more abundant in the Olfm4-L neutrophils. The data suggest different preparedness to infection in the subsets. In the limited material analysed here, there was no significant correlation between the severity of sepsis and the proportion of Olfm4-H neutrophils, but an increased concentration of Olfm4 in plasma from septic shock patients as compared to healthy blood donors was observed. Furthermore, in neutrophil subsets isolated from septic shock patients, 28 proteins had significantly higher abundance in the Olfm4-H subset and 38 in the Olfm4-L subset, the latter including *e.g.* Fc receptor proteins and MHC class I molecules, suggesting distinct immunological responses. This is the first report pointing towards differential functions of the Olfm4-defined neutrophil subpopulations in humans and the data are consistent with the idea of distinct responses in the subsets during infection and inflammation.

## Introduction

The neutrophil granulocyte, the most abundant of the leukocytes, is today recognized as a heterogenous cell type with distinct subsets [1]. The glycoprotein olfactomedin-4 (Olfm4) marks such a subset in humans as well as in mice [2–5]. Olfm4 is a constitutive human neutrophil subset marker, meaning that the Olfm4 protein is present in only a subset of neutrophils in the peripheral blood of a given individual [2, 3, 5], independently of cell activation status or age [3].

The Olfm4 protein locates to the specific granules of neutrophils [2, 3]. All granulocyte precursors express Olfm4 mRNA at the myelocyte and metamyelocyte stages, indicating that an unknown post-transcriptional mechanism determines whether the protein is translated [2]. The proportion of Olfm4-high (Olfm4-H) neutrophils varies between individuals (1-70% of neutrophils) [2–4, 6] and healthy blood donors display low variation in the proportion of Olfm4-H neutrophils over time [3]. Meanwhile, proportional fluctuations over time have been observed in children after bone marrow transplantation [7], and an increase in the Olfm4-H proportion has been observed following haemorrhagic shock [8]. The function of human Olfm4 and potential differential functions of the subset are not known.

We have previously shown that the Olfm4-defined neutrophil subsets have equal tendency to phagocytose bacteria, undergo apoptosis, and transmigrate to inflamed tissue, and that Olfm4 can be found in a subset of neutrophil extracellular traps (NETs) [3]. It has more recently been discovered that the proportion of Olfm4-H neutrophils correlates with the severity of paediatric septic shock [4] and can independently predict mortality in adult septic shock [6]. Earlier mouse studies suggested a detrimental effect of Olfm4 during bacterial infection, with mice lacking Olfm4 having an enhanced capacity to combat *Staphylococcus aureus* and *Escherichia coli* as well as prevent colonization with *Helicobacter pylori* [9]. In a cancer context, Olfm4 has been shown to have tissue-specific tumor-promoting or tumor-suppressive activity [9].

Sepsis is the leading cause of death among hospitalized patients worldwide with a mortality rate of ~20% [10]. According to the consensus definition Sepsis-III [11], sepsis is life-threatening organ dysfunction caused by a dysregulated host response to infection. Septic shock is a subset of sepsis in which the underlying circulatory and cellular metabolism abnormalities are profound enough to substantially increase mortality. The septic immune response is multifaceted with components of exaggerated activation and immune suppression, and the response varies between individuals [10]. During the septic immune response, the innate immune system is severely dysregulated. Neutrophil granulocytes contribute to microbe clearance but are also inappropriately activated leading to immunopathology [12, 10]. Consequently, the finding that the Olfm4-H neutrophil proportion correlates with the severity and outcome of sepsis [4, 6] warrants investigation into the differential contributions of the subsets to immune dysfunction leading to sepsis pathogenesis.

The aim of this study was to decipher proteomic differences between the Olfm4-defined human neutrophil subsets at baseline and in the context of septic shock. The hypothesis was that Olfm4 marks a neutrophil subset with a distinct proteomic profile, predisposing it for detrimental processes in sepsis.

## Materials and Methods

### Patient characteristics and data collection

Adult patients (>18 years old) admitted to the Intensive Care Unit (ICU) at Linköping University Hospital with a diagnosis of septic shock using the Sepsis-III criteria were included within 12 h of admission. Acute coronary syndrome at presentation was an exclusion criterion. The patients were part of a larger on-going, prospective study (NCT04695119). The mean age of the 20 patients included for analysis of Olfm4-H proportion and plasma Olfm4 concentration was 67 years (range 46-85); 12 were males and 8 females. Data on sequential organ failure assessment (SOFA) score on admission and number of days alive and free of invasive ventilatory, renal or hemodynamic support within 30 days, termed organ support-free days (OSFD), were collected. The SOFA score is the predominant score quantifying the severity of organ dysfunctions in sepsis [11], and OSFD is a commonly used proxy of outcome in septic shock studies [13]. The outcome measures were the correlation between the percentage of Olfm4-H neutrophils and SOFA score or OSFD, and between the plasma concentration of Olfm4 and SOFA score or OSFD.

### Neutrophil isolation from peripheral blood

Granulocytes were isolated from whole blood of healthy blood donors (18 ml) through the Linköping University Hospital blood bank or septic shock patients (see above, 3 ml) within three hours of blood collection, by density gradient centrifugation. The blood was drawn in EDTA BD Vacutainer blood collection tubes (Becton Dickinson, Franklin Lakes, NJ, USA) and layered onto a gradient of Polymorphprep (Alere Technologies, Oslo, Norway) and Lymphoprep (Alere Technologies) according to the manufacturer’s instructions. Briefly, tubes were then centrifugated in a swing-out centrifuge for 40 minutes at 480 × *g* at room temperature (RT), and the granulocyte band was washed in phosphate-buffered saline (PBS) at RT. Remaining erythrocytes were lysed with distilled H_2_O at 4°C and the granulocytes washed twice with Krebs-Ringer glucose (KRG) phosphate buffer at 4°C, resuspended in PBS on ice and processed immediately.

### Antibody staining and FACS

Granulocytes were stained for surface CD15 followed by intracellular Olfm4. Briefly, 2×10^7^ cells resuspended in 0.5 ml PBS were incubated with PE-conjugated mouse anti-human CD15 antibody (BioLegend, San Diego, CA, USA) diluted 1:40, for 30 minutes on ice. Cells were then washed in PBS and resuspended in 1 ml cold BD Cytofix/Cytoperm (Becton Dickinson) for fixation and permeabilization, incubating for 20 minutes on ice. After washing twice with cold BD Perm/Wash (Becton Dickinson), the cells were resuspended in 1 ml BD Perm/Wash containing rabbit anti-Olfm4 serum (prepared in-house and kindly provided by Dr. Matthew Alder, Cincinnati Children’s Hospital, OH, USA [7]) diluted 1:400, and incubated for 30 minutes at RT. The cells were washed twice and resuspended in 1 ml BD Perm/Wash containing Alexa Fluor 647-conjugated goat anti-rabbit IgG (Thermo Fisher Scientific, Waltham, MA, USA) diluted 1:400, incubating for 30 minutes at RT. After three final washing steps, the pellet was resuspended in 300 μl PBS for fluorescence-activated cell sorting (FACS).

The fixed and stained granulocytes were sorted into two populations; CD15-high and Olfm4-high (Olfm4-H) or CD15-high and Olfm4-low (Olfm4-L) neutrophils, on a BD FACS Aria III Cell Sorter (Becton Dickinson). Between 1×10^6^ and 5.5×10^6^ cells per subset were sorted, depending on the subset proportions in different individuals. The gating strategy is shown in supplementary Fig. S1.

### Peptide preparation and LC-MS/MS

A protocol for the preparation of formaldehyde-fixed tissue for subsequent liquid chromatography with tandem mass spectrometry (LC-MS/MS) analysis published by Coscia *et al*. [14] was adapted to formaldehyde-fixed neutrophils in suspension. Sorted neutrophils were pelleted and resuspended in 100 μl lysis buffer (300 mM Tris/HCl in 50% acetonitrile, pH 8). Samples were sonicated using a Fisherbrand model 705 sonic dismembrator (Thermo Fisher Scientific) for 2 cycles (5 seconds on/off) at 40 amplitude and placed in a heating block for 90 minutes at 90° C. To reduce proteins, samples were incubated in 5 mM 1,4-dithiothreitol (Roche, Basel, Switzerland) for 20 minutes, mixing at 1400 RPM, at RT. Then, to alkylate proteins, 25 mM iodoacetamide (Merck, Darmstadt, Germany) was added and samples incubated for a further 30 minutes, mixing at 1400 RPM, at RT in the dark. Protein samples were vacuum concentrated to about 20 μl using a CHRIST RVC 2-25 centrifuge (Martin Christ Gefriertrocknungsanlagen, Osterode am Harz, Germany) set to 60° C. The protein concentration was measured by Nanodrop (Thermo Fisher Scientific) at 280 nm, and 100 μg of protein was digested by adding MS grade Pierce Trypsin Protease (Thermo Fisher Scientific) at a protein/trypsin ratio of 1:50 and 10 % acetonitrile (Thermo Fisher Scientific), in a final volume of 100 μl. Samples were incubated overnight, shaking at 1400 RPM, at 37° C.

Peptide samples were acidified by adding 1 % trifluoroacetic acid (Merck) and debris pelleted by centrifugation for 5 minutes at 14 000 × *g*. Peptide clean-up was performed using Pierce C18 tips (Thermo Fisher Scientific) following the manufacturer’s instructions, eluting using 80 % acetonitrile (Thermo Fisher Scientific). Eluted samples were dried completely by CHRIST RVC 2-25 centrifuge at 45° C and resuspended in 15 μl 0.1 % formic acid (Sigma-Aldrich, Saint Louis MO USA), vortexed and sonicated in an ultrasonic water bath for 10 minutes. After a further concentration measurement, samples were diluted to 0.1 μg/μl in 0.1 % formic acid in MS autosampler vials (Thermo Fisher Scientific). Five μl of peptides were loaded onto an Easy nanoLC (Thermo Fisher Scientific) coupled to a Q Exactive HF Hybrid Quadrupole-Orbitrap Mass Spectrometer (Thermo Fisher Scientific). Peptides were separated by reverse phase chromatography using a C18 pre-column (Acclaim PepMap 100, 75μm×2cm, Thermo Fisher Scientific) followed by C18 reversed-phase LC column (PepMap RSLC C18, 2μm, 100A 75μm×25cm, Thermo Fisher Scientific) on EASY nLC 1200 system (Thermo Fisher Scientific). A linear gradient from 0.1% formic acid in H_2_O (A) to 0.1% formic acid in 80% acetonitrile (B) was applied at a flow rate of 300 nL/min as follows: from 6 % B to 28 % B at 50 minutes; from 28 % B to 40 % B at 78 minutes, 100% B at 95 minutes. Measurements were taken in positive polarity, Full MS to dd-MS^2^ mode. Resolution was 120 000 with a range of 380 to 1400 m/z and an automatic gain control target of 3×10^6^ and 120 ms ion time.

### Bioinformatics

Obtained RAW-files were analyzed in Proteome Discoverer 2.5.0.4 (Thermo Fisher Scientific) using a human reference proteome database (79038 entries, downloaded from Uniprot 220215). MSPepSearch and Sequest HT were searched in sequence, keeping medium confidence peptides between, with fragment mass tolerance set to 0.15 Da and precursor mass tolerance set to 14 PPM. Full trypsin digestion and a maximum of two missed cleavages as well as a minimum peptide length of 6 amino acids was specified. Carbamidomethyl of cysteine was specified as a fixed modification and oxidation as dynamic modification. Percolator validation used standard settings. Label free quantification was carried out to calculate abundances of proteins in Olfm4-H and Olfm4-L neutrophil samples. Unique and razor peptides were considered in protein quantification. Precursor abundance was based on intensity. To account for experimental bias, normalization of protein abundances were based on a set of proteins previously described as “housekeeping proteins” [15] and imputation was set to replicate based resampling. Peptides of less than medium confidence were filtered out and proteins were identified by at least 2 peptides. Protein abundances were calculated by summing sample abundances of connected peptide groups. Median paired Olfm4-H/L log2 ratio calculations were based on all possible pairwise ratios of connected peptides between replicates and tested for significant difference by a background-based t-test with Benjamini-Hochberg (BH) correction. Proteome Discoverer filters were set to include high confidence master proteins with an abundance value in both Olfm4-H and Olfm4-L groups. Proteins in the Olfm4-H and Olfm4-L samples where the log2 abundance ratio had a *p-*value of < 0.1 were considered and the corresponding gene symbols used as input for pathway enrichment analysis in the Reactome pathway database (https://reactome.org/; release 79) [16] with regards to the human genome. Reactome is free, open-source, open-data and peer-reviewed knowledgebase of biomolecular pathways providing high-quality curated data over other open-source databases [17]. A minimum of 5 genes were considered to calculate significantly enriched pathways with BH-corrected *p*-value < 0.05.

### Imaging flow cytometry of Olfm4-defined subset proportions

Whole blood from septic shock patients (3 ml) was collected in EDTA BD Vacutainer blood collection tubes. The patient samples were processed as soon as possible, generally within three hours, but never later than 20 h after blood was drawn. Control experiments where blood from 3 healthy donors was processed immediately upon arrival or incubated for 20 h at RT before processing showed no impact of incubation on the percentage of Olfm4-H neutrophils (supplementary Fig. S2a). To fix leukocytes and remove erythrocytes, 400 μl of blood was treated with BD FACS lysing solution (Becton and Dickinson) according to the manufacturer’s instructions. Plasma was separated from the remaining blood volume and frozen at −70°C for later analysis of free Olfm4 by enzyme-linked immunosorbent assay (ELISA) as described below. The fixed leukocytes were stained for the neutrophil marker CD15 followed by permeabilization and staining of Olfm4 as described above, with the addition of DAPI at 60 nM in the secondary antibody step to stain DNA. Each patient sample was stained in parallel with a sample from an anonymous healthy blood donor donating blood to the Linköping University Hospital blood bank. The cells were resuspended in 20 μl PBS for imaging flow cytometry. The size of the Olfm4-H neutrophil subset was determined by using an ImageStream MK II (Luminex corporation, Austin, TX, USA). At least 1500 and up to 10 000 events were acquired at 60X magnification. IDEAS version 6.2 was used for analysis. After applying a colour compensation matrix, neutrophils were identified by gating for focus followed by aspect ratio and size, DAPI intensity, CD15 intensity and finally the Olfm4-H and Olfm4-L populations were identified based on Olfm4 intensity (supplementary Fig. S3).

### ELISA

After centrifugation of the thawed plasma samples at 2000 × *g* for 10 minutes and dilution of samples by 1:10 or 1:20, the ab267805 Human Olfm4 SimpleStep ELISA kit (Abcam, Cambridge United Kingdom) was performed in duplicate according to the manufacturer’s instructions and Olfm4 concentrations in plasma were calculated using a linear standard curve. The mean Olfm4 concentration of the duplicates is reported. In addition, the Olfm4 concentration was normalized to the neutrophil concentration in the original blood sample, as estimated from imaging flow cytometry sample information.

Since some patient samples were processed the morning after collection (see above), we examined whether incubation of the blood affected the concentration of plasma Olfm4. Blood from 3 healthy donors was processed immediately upon arrival or incubated for 20 h at RT before processing. Incubation did not significantly affect the concentration of Olfm4 in plasma although there was a trend toward higher concentrations upon incubation (supplementary Fig. S2b).

### Statistical methods

Statistical analysis was performed using GraphPad Prism, v. 9.2.0. See above for statistical analysis of proteomics data. All box-and-whisker plots depict median and interquartile range. Differences between groups were analysed by the Mann-Whitney U test unless otherwise stated and the result considered significant at *p*<0.05. Throughout the manuscript significant differences are denoted by * (*p*<0.05), ** (*p*<0.01), or *** (*p*<0.001).

## Results

### The Olfm4-defined neutrophil subsets from healthy donors exhibit distinct proteomic profiles

The Olfm4-H neutrophil subset has been proposed to play a pathogenic role in septic shock [4, 6] and the hypothesis of this work was that Olfm4 marks a neutrophil subset with a distinct proteomic profile, predisposing it for detrimental processes in sepsis. To determine whether the two subsets exhibit differential protein production at baseline, the proteomes of FACS-sorted Olfm4-H and Olfm4-L resting neutrophils from healthy blood donors were analysed. A published method [14] was adapted to suit neutrophils in suspension, enabling analysis of formaldehyde-fixed, permeabilized and antibody-stained neutrophils by LC-MS/MS. The approach yielded a similar number of proteins and peptide groups identified in both subsets (Fig 1a). In total, 1136 protein were identified in both Olfm4-H and Olfm4-L neutrophil samples. Supplementary Table S1 lists all the identified proteins and their abundance in each subset. Among the 20 proteins with the highest abundance, major neutrophil granule proteins such as lysozyme, myeloblastin, S100A8/A9, azurocidin, neutrophil elastase, cathepsin G, lactoferrin, and myeloperoxidase were found, and none of these displayed significantly increased abundance in either subset (Fig. 1b). Olfm4 was detected among the 20 most abundant proteins in the Olfm4-H subset, and although there was a clear difference in abundance of Olfm4 between the subsets with a nearly 24-fold increase in the Olfm4-H subset, Olfm4 was detected in both populations (Supplementary Table S1). Thus, we suggest using the terms “Olfm4-high” and “Olfm4-low” to describe the subsets rather than “Olfm4-positive” and “Olfm4-negative.” The relative abundance of each protein between the Olfm4-H and Olfm4-L subsets was analysed in terms of log2 abundance ratio, where a positive value means a higher abundance in the Olfm4-H subset and vice versa. Olfm4 had a median log2 ratio between the Olfm4-H and Olfm4-L subsets of 4.1, with a significance level of *p*=8.0×10^−16^ (Fig. 1b).

**Figure 1.**
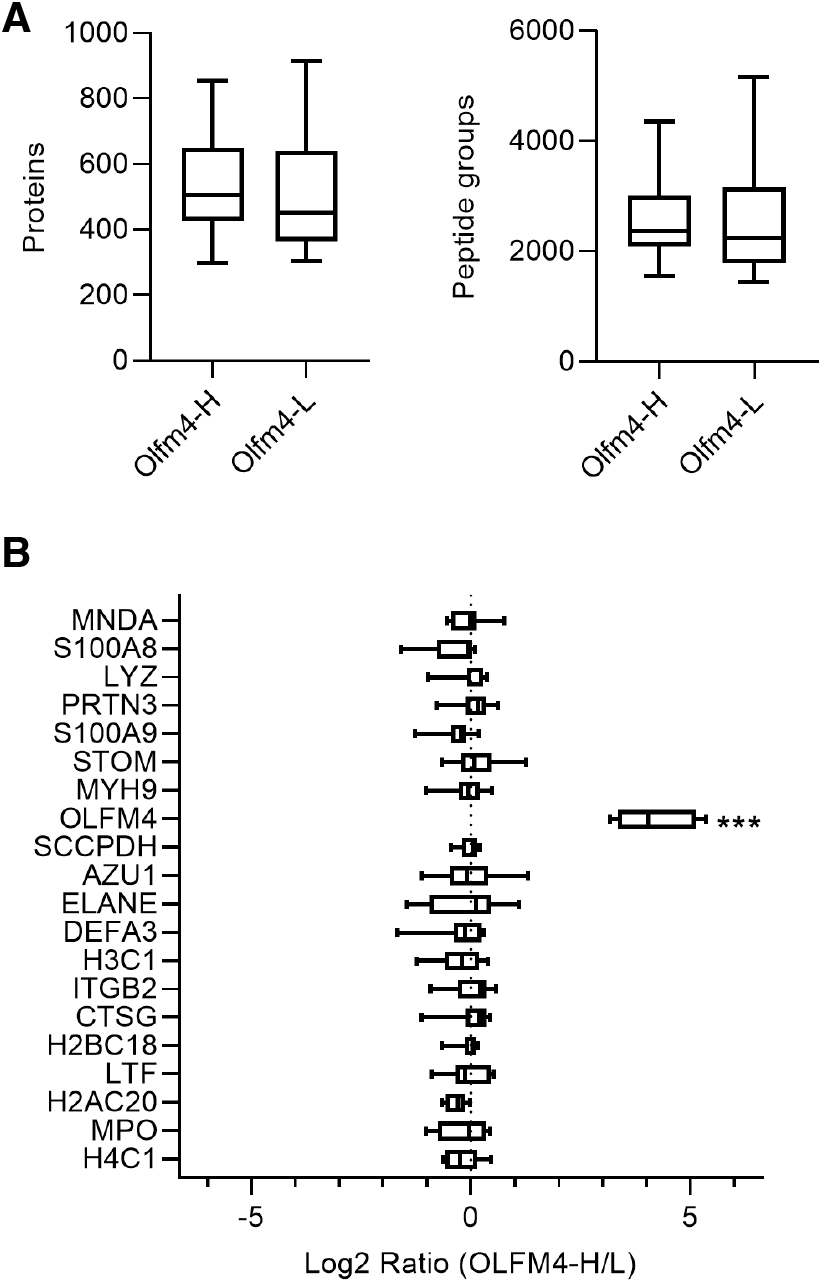
Proteomic analysis of Olfm4-defined neutrophil subsets in healthy blood donors. Olfm4-H and Olfm4-L neutrophils isolated from healthy blood donors (n=8) were sorted by FACS and analysed by LC-MS/MS. (A) Box-and-whisker plots showing the number of proteins and peptide groups identified in Olfm4-H and Olfm4-L neutrophil samples. (B) Box-and-whisker plot showing protein abundance log2 ratios for the 20 most abundant proteins in paired Olfm4-H/L neutrophil samples. The probable contaminant desmoplakin has been omitted.

A log2 abundance ratio between the Olfm4-H and Olfm4-L subsets with a *p-*value of < 0.1 was observed for 110 proteins, where 47 proteins were more abundant in the Olfm4-H and 62 in the Olfm4-L subset (Fig. 2a). In search of a surrogate marker for Olfm4, it was of interest to determine whether any other protein displayed a bimodal distribution, meaning that for all eight analysed blood donors, the log2 abundance ratio was either only positive or only negative. In addition to Olfm4, 27 of the differently abundant proteins displayed such a bimodal distribution (Supplementary Table S1). Several proteins relevant to the neutrophil immune response such as Rab3D, Rab11A, leucine rich alpha-2 glycoprotein, cytochrome b-245 chaperone 1 (CYBC1) and V-type proton ATPase 16 kDa subunit showed higher abundance in the Olfm4-H subset (Fig. 2b). Meanwhile, S100-A7, Rab3A, C-X-C chemokine receptor type 1 (CXCR1), and neutrophil defensin alpha 4 among others displayed higher abundance in the Olfm4-L subset (Fig 2c).

**Figure 2.**
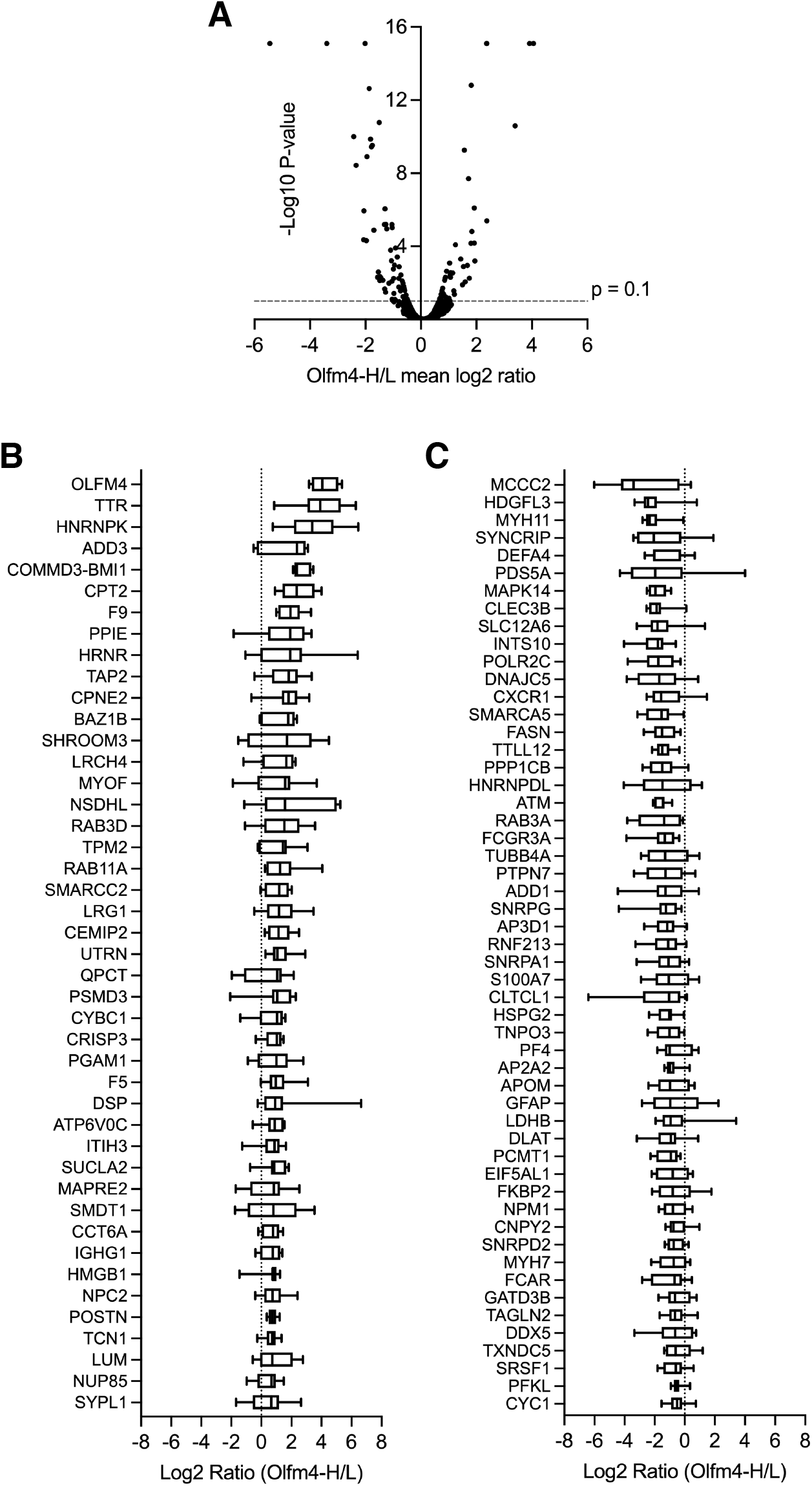
Differential proteomic profiles of Olfm4-H and Olfm4-L neutrophil subsets in healthy blood donors. Olfm4-H and Olfm4-L neutrophils isolated from healthy blood donors (n=8) were sorted by FACS and analysed by LC-MS/MS. (A) Volcano plot showing median log2 abundance ratios between the identified proteins in Olfm4-H and Olfm4-L neutrophils, and their *p-*values. (B) Box-and-whisker plot showing the 47 proteins with a positive log2 abundance ratio and p < 0.1, indicating increased abundance in the Olfm4-H neutrophils. (C) Box-and-whisker plot showing the 62 proteins with a negative log2 abundance ratio and p < 0.1, indicating increased abundance in the Olfm4-L neutrophils.

The list of differentially abundant proteins converted to gene symbols was used as the input list to the Reactome database for pathway enrichment analysis. In healthy blood donors, the Olfm4-H subset showed significant enrichment in the pathways neutrophil degranulation (*p=*1.4×10^−8^), innate immune system (*p*=3.6×10^−4^) and immune system (*p*=3.8×10^−2^) (Fig. 3a, Supplementary table S2a), and the Olfm4-L subset in the neutrophil degranulation pathway (*p*=1.3×10^−2^) only (Fig. 3b; Supplementary table S2b).

**Figure 3.**
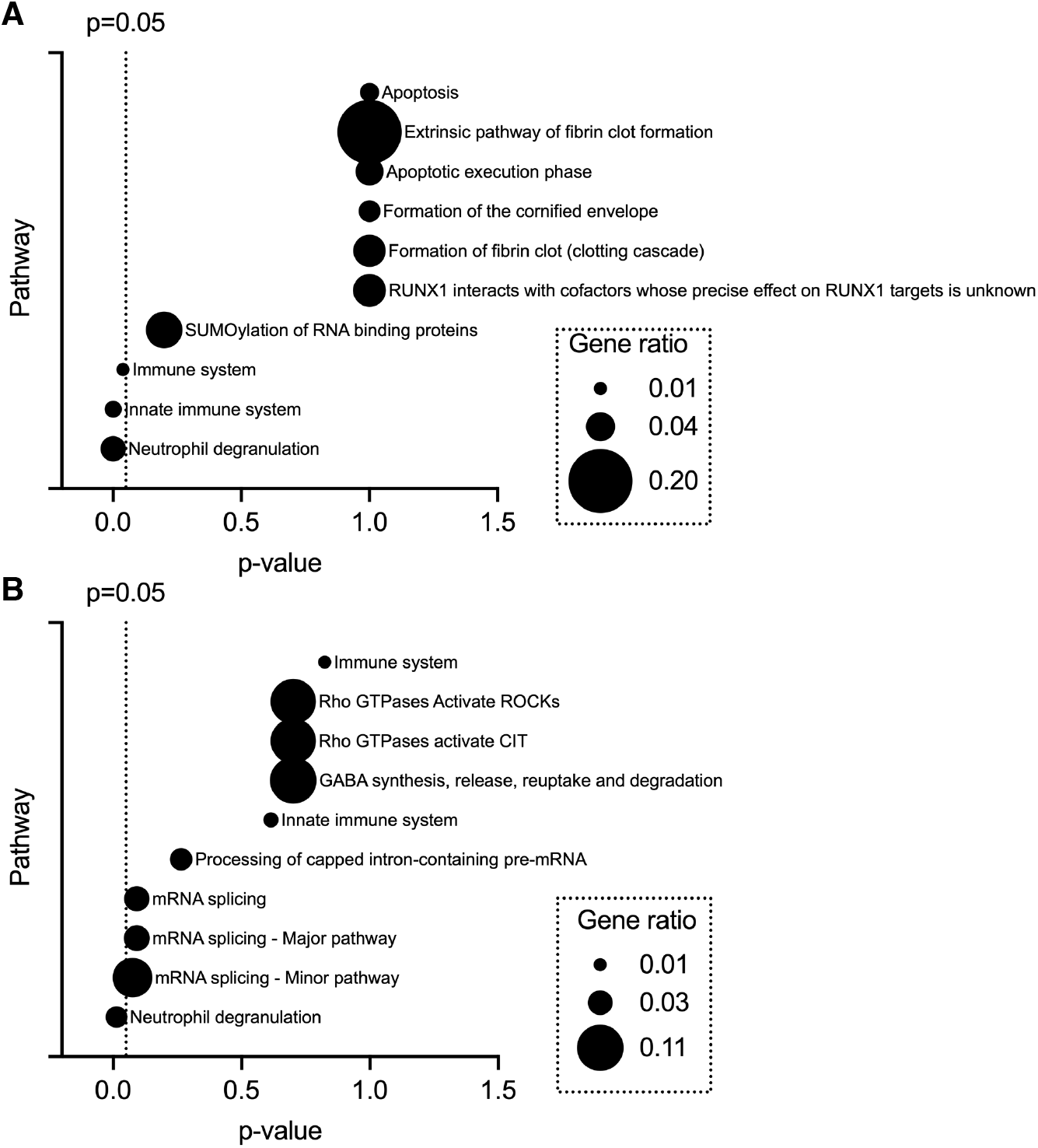
Reactome pathway analysis of differentially abundant proteins in the Olfm4-defined subsets in healthy blood donors. Gene symbols for the identified proteins with a log2 abundance ratio between the Olfm4-H and Olfm4-L subsets from healthy blood donors with a *p*-value of < 0.1 were analysed by pathway enrichment analysis. The gene ratio was calculated by dividing the number of input genes by the number of genes in the database for each enriched pathway. (A) Diagram showing the ten most enriched pathways in the Olfm4-H neutrophil subset. (B) Diagram showing the ten most enriched pathways in the Olfm4-L neutrophil subset. The x-axes show BH-corrected *p*-values.

### Olfm4-defined subset proportions and plasma Olfm4 in septic shock

Having discovered that Olfm4-H and Olfm4-L neutrophils display different proteomic profiles at baseline and as the literature suggests a pathogenic role for the Olfm4-H neutrophils in sepsis [4, 6, 18], we next focused on the role of the Olfm4-defined neutrophil subsets in septic shock. First, we sought to corroborate the correlation between the percentage of Olfm4-H neutrophils and disease severity in an adult population of septic shock patients admitted to the ICU at Linköping University Hospital. The Olfm4-proportions in whole blood neutrophils, as well as plasma Olfm4 concentrations, were analysed in 20 patients and 20 healthy blood donors. Data on SOFA score [11] on admission, where a higher score corresponds to a higher degree of organ dysfunction, and OSFD for the first 30 days [13], where a higher value means that the patient was alive and free of invasive organ support for a greater number of days during the first month, were collected. The mean admission SOFA score was 7.9 (median 8.5, range 1-15) and the mean OSFD was 20.4 (median 26, range 0-29).

The proportion of Olfm4-H neutrophils was similar between septic shock patients and healthy blood donors (Fig. 4a). The percentage of Olfm4-H neutrophils showed a non-significant correlation with SOFA score (Fig. 4b left panel) and no tendency for correlation with OSFD (Fig. 4b, right panel). As the percentage of Olfm4-H neutrophils has previously been shown to predict mortality in septic shock, the patients were divided into those with a low versus high proportion of Olfm4-H neutrophils using the previously published cut-off of >37.6% Olfm4-H neutrophils [6]. Patients with a high proportion tended to have higher SOFA scores upon admission to the ICU, but the difference was not statistically significant (Fig. 4c, left panel) and there was no trend regarding OSFD (Fig 4c, right panel). The concentration of plasma Olfm4 was drastically augmented in septic shock patients as compared to healthy controls (Fig. 4d, left panel). However, neutrophilia is a common finding in sepsis [19], and when the plasma Olfm4 values were normalised to the estimated number of neutrophils in the sample (based on the neutrophil concentration as recorded by imaging flow cytometry), there was no longer a significant difference between patients and controls, although the trend remained (Fig. 4d, right panel). The plasma Olfm4 concentration showed no correlation with either SOFA score or OSFD (data not shown). In summary, in the limited material analysed here the percentage of Olfm4-H neutrophils did not correlate significantly with the severity of septic shock, although there was such a trend. Meanwhile, septic shock patients had drastically elevated concentrations of free Olfm4 in plasma, associated with neutrophilia.

**Figure 4.**
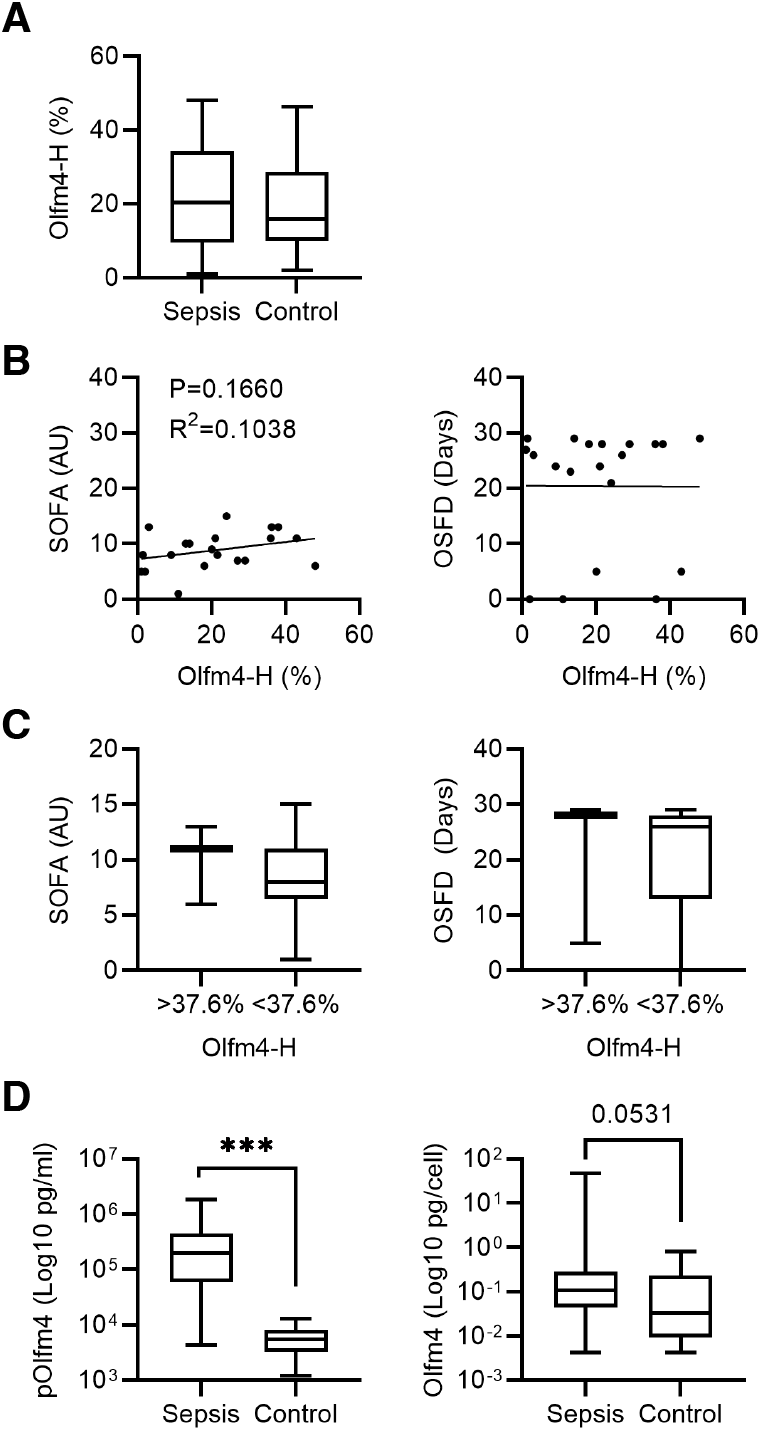
Analysis of Olfm4-defined neutrophil subset proportions and plasma Olfm4 concentrations in septic shock patients. Peripheral blood was collected from septic shock patients (n=20) and healthy blood donors (n=20). The proportion of Olfm4-H neutrophils was analysed by imaging flow cytometry after fixation of leukocytes, staining of CD15 for neutrophil identification, permeabilization and antibody staining of Olfm4. ELISA was used to analyse Olfm4 concentrations in plasma. (A) Box-and-whisker plot showing the proportion of OLFM4-H neutrophils in sepsis patients and healthy controls. (B) Scatterplots showing the correlation between the proportion of Olfm4-H neutrophils and SOFA score (left panel) or OSFD (right panel). (C) Box-and-whisker plots showing SOFA score (left panel) or OSFD (right panel) of sepsis patients with percentage of Olfm4-H neutrophils above 37.6% (n=3) or below 37.6% (n=17). (D) Box-and-whisker plots showing raw plasma Olfm4 concentrations (left panel) and adjusted to the estimated concentration of neutrophils in the sample (right panel).

### The Olfm4-defined neutrophil subsets exhibit proteomic differences in septic shock

The data above revealed that the Olfm4-defined subsets display different proteomic profiles at baseline, and the literature suggests a pathogenic role for Olfm4-H neutrophils in sepsis. Accordingly, we hypothesized that the Olfm4-H neutrophils display a distinct proteomic profile also in septic shock, disposing them to detrimental processes.

The proteomes of FACS-sorted Olfm4-H and Olfm4-L resting neutrophils from three septic shock patients were individually analysed, using the same workflow as for healthy blood donors described above. A similar number of proteins and peptide groups were identified in both subsets (Fig 5a). The total number of proteins found was 1116, which was very similar to what was found for blood donors. Supplementary table S3 lists all the identified proteins and their abundance in each subset in the patients. The most abundant proteins included S100A8/A9, azurocidin, neutrophil elastase, cathepsin G, lactoferrin, and myeloperoxidase (Fig. 5b). These proteins are comparable to the most abundant proteins in the blood donor neutrophils, and none of the most abundant proteins apart from Olfm4 displayed a significantly increased abundance in either of the subsets. Olfm4 was again among the 20 most abundant proteins in the Olfm4-H subset and had a median log2 ratio between the Olfm4-H and Olfm4-L subsets of 3.74, with a significance level of *p=*4×10^−15^ (Fig. 5b).

**Figure 5.**
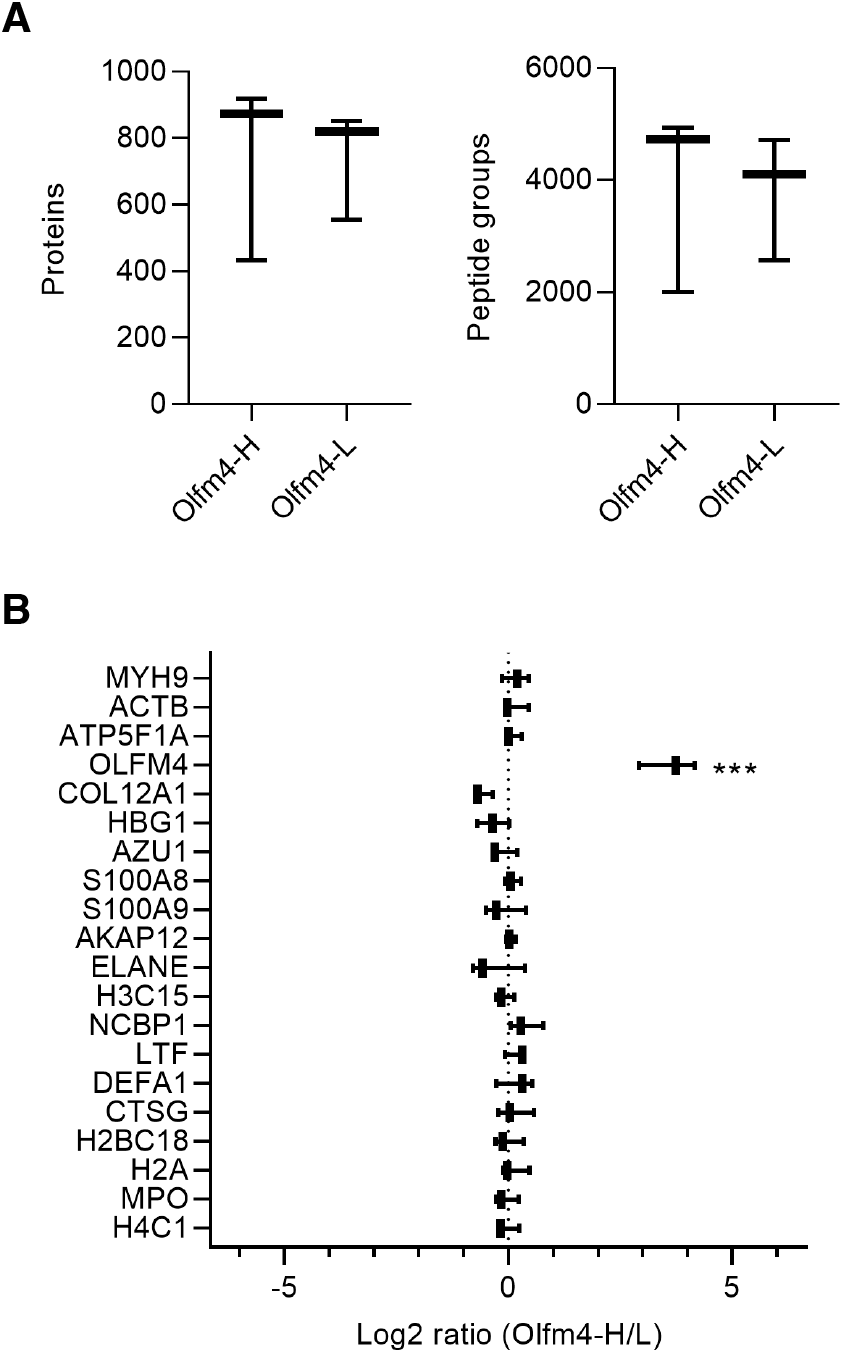
Proteomic analysis of Olfm4-defined neutrophil subsets in septic shock patients. Olfm4-H and Olfm4-L neutrophils isolated from septic shock patients (n=3) were sorted by FACS and analysed by LC-MS/MS. (A) Box-and-whisker plots showing the number of proteins and peptide groups identified in Olfm4-H and Olfm4-L neutrophil samples. (B) Box-and-whisker plot showing protein abundance log2 ratios for the 20 most abundant proteins in paired Olfm4-H/low neutrophil samples.

A log2 abundance ratio between the Olfm4-H and Olfm4-L subsets with a *p-*value of < 0.1 was seen for 66 proteins, where 28 proteins were more abundant in the Olfm4-H and 38 in the Olfm4-L subset (Fig. 6a). In addition to Olfm4, 47 other proteins displayed a bimodal distribution (Supplementary Table S3). Proteins such as nod-like receptor family member X1 and CYBC1 displayed higher abundance in the Olfm4-H subset (Fig. 6b) and among others Fc epsilon receptor 1 gamma subunit, Fc gamma receptor IIa, Wiskott Aldrich Syndrome protein and scar homologue (WASH) complex subunit 4, and human leukocyte antigen (HLA) class I histocompatibility antigen A alpha chain in the Olfm4-L subset (Fig 6c).

**Figure 6.**
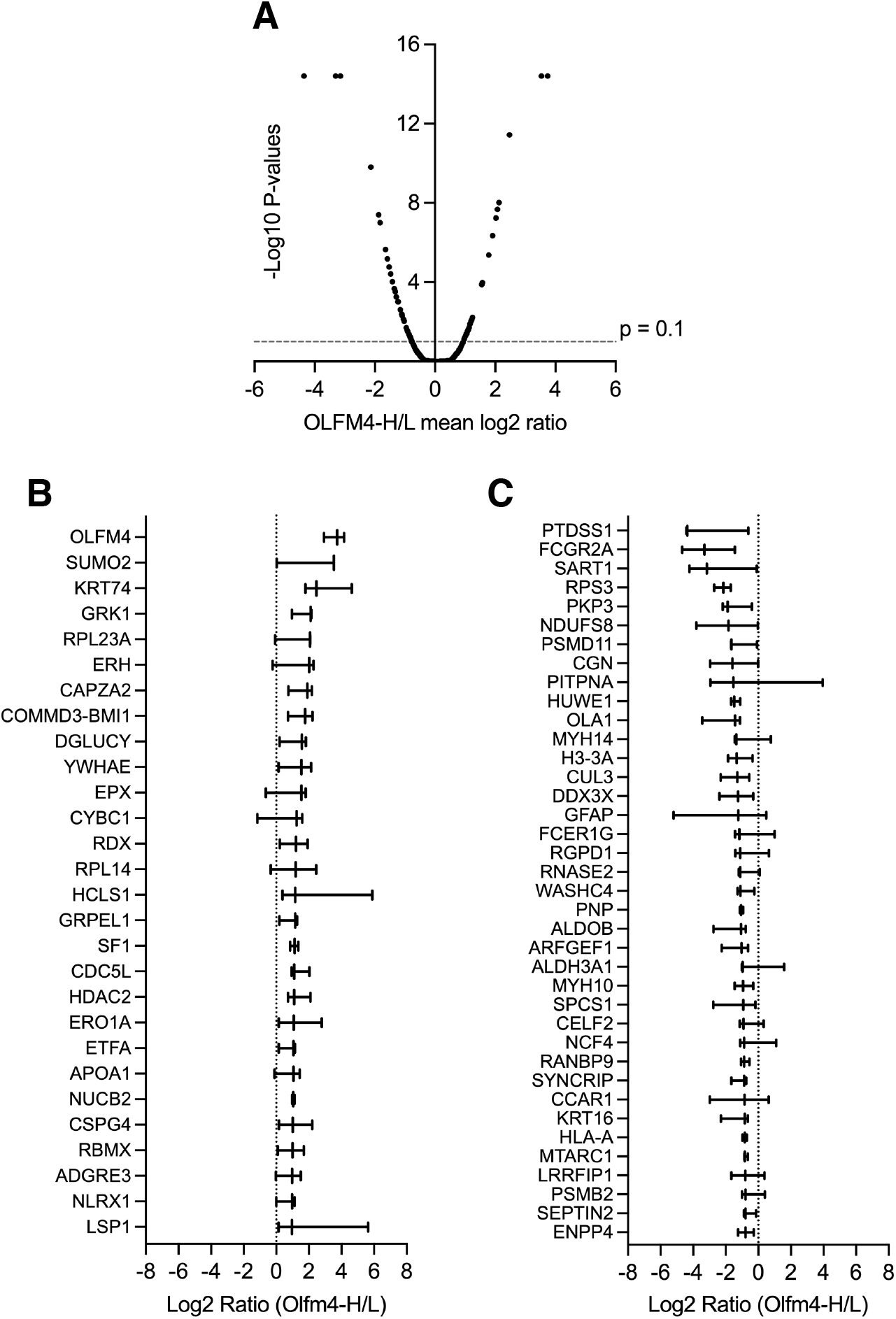
Differential proteomic profiles of Olfm4-H and Olfm4-L neutrophil subsets in septic shock patients. Olfm4-H and Olfm4-L neutrophils isolated from septic shock patients (n=3) were sorted by FACS and analysed by LC-MS/MS. (A) Volcano plot showing median log2 abundance ratios between the identified proteins in Olfm4-H and Olfm4-L neutrophils, and their *p-*values. (B) Box-and-whisker plot showing the 28 proteins with a positive log2 abundance ratio and p < 0.1, indicating increased abundance in the Olfm4-H neutrophils. (C) Box-and-whisker plot showing the 38 proteins with a negative log2 abundance ratio and p < 0.1, indicating increased abundance in the Olfm4-L neutrophils.

Pathway enrichment analysis showed no statistically significant enrichment in any pathway in either the Olfm4-H (Fig. 7a, supplementary table S4a) or Olfm4-L (Fig. 7b, supplementary table S4b) subset in the septic shock patients. However, the Olfm4-L subset showed near-significant enrichment in the pathways neutrophil degranulation (*p*=7.1×10^−2^), innate immune system (*p*=7.3×10^−2^) and antigen processing – cross presentation (*p*=9.5×10^−2^).

**Figure 7.**
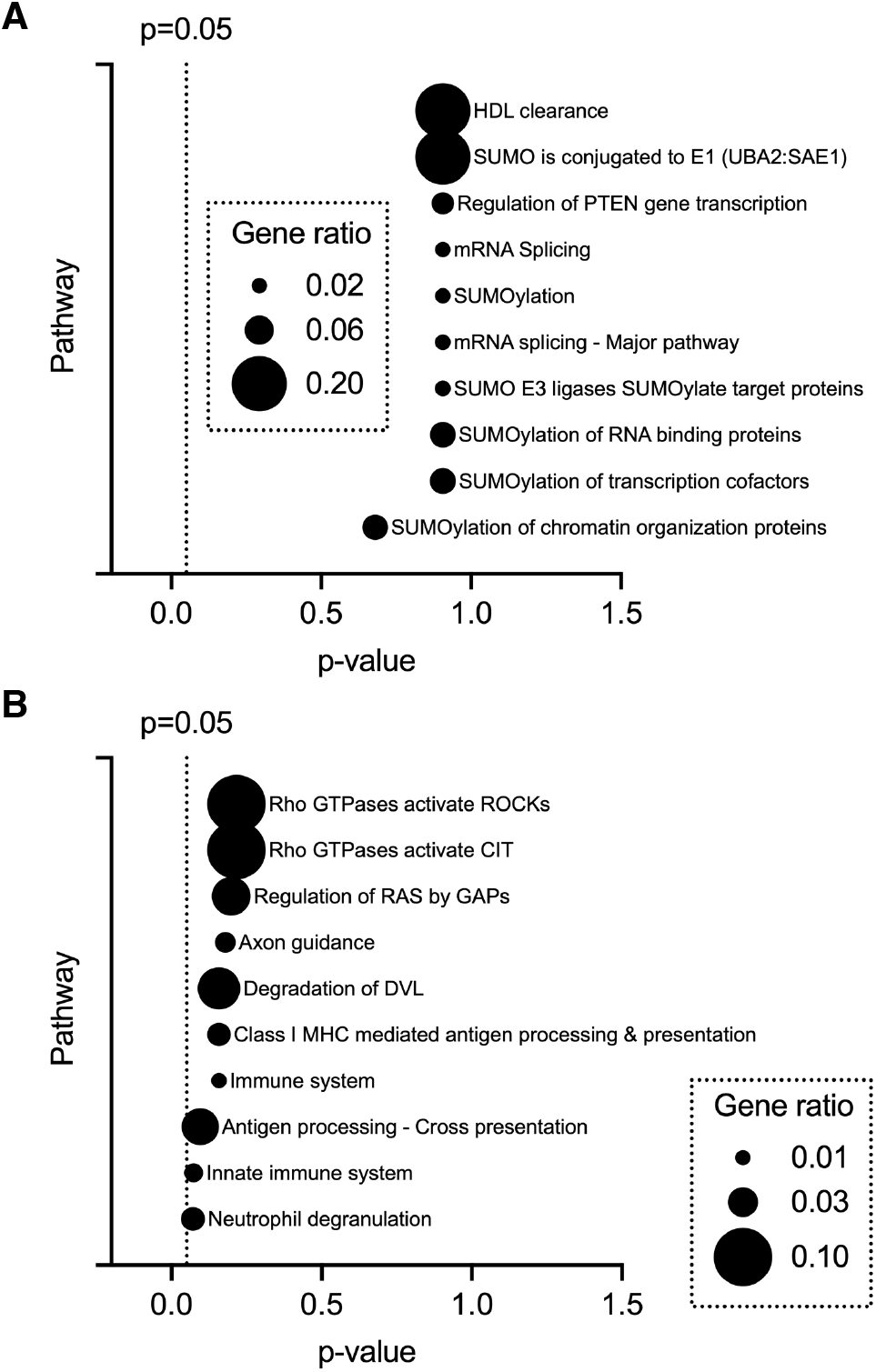
Reactome pathway analysis of differentially abundant proteins in the Olfm4-defined subsets in septic shock patients. Gene symbols for the identified proteins with a log2 abundance ratio between the Olfm4-H and Olfm4-L subsets in septic shock patients with a *p-*value of < 0.1 were analyzed by pathway enrichment analysis. The gene ratio was calculated by dividing the number of input genes by the number of genes in the database for each enriched pathway. (A) Diagram showing the ten most enriched pathways in the Olfm4-H neutrophil subset. (B) Diagram showing the ten most enriched pathways in the Olfm4-L neutrophil subset. The x-axes show BH-corrected *p*-values.

## Discussion and conclusion

The proportion of Olfm4-H neutrophils has been found to correlate with the severity of septic shock [4, 6], but functional differences between the Olfm4-defined subsets in sepsis are unknown. To approach the hypothesis that Olfm4 marks a subset with a proteomic profile that predisposes it to detrimental immunological processes in sepsis, we deciphered the proteomic differences between the Olfm4-defined human neutrophil subsets in resting neutrophils as well as in neutrophils from septic shock patients. Thus, we revealed differences between the subsets in proteins important for the neutrophil immune response.

This is to our knowledge the first report of the proteome of fixed, permeabilized, antibody-stained and FACS-sorted neutrophils, and a protocol developed for formaldehyde-fixed tissue sections [14] was adapted for fixed cells in suspension. In the proteomic analysis of sorted Olfm4-defined neutrophil subsets from healthy blood donors, 1136 proteins were identified, which is only slightly lower than a previously published proteomics study of non-fixed human neutrophil granules that identified about 1500 proteins [20]. Thus, the new method had satisfactory sensitivity. The analysis showed that similar numbers of proteins were identified in the subsets, and major neutrophil granule proteins were present in equal abundance in both subsets, as previously published for e.g. neutrophil gelatinase-associated lipocalin (NGAL) [3] and gelatinase [2]. The only exception was the Olfm4 protein itself, which as expected was vastly more abundant in the Olfm4-H subset. The finding that the Olfm4 protein was detected in the Olfm4-L subset, albeit at a 24 times lower level than in the Olfm4-H subset, is in accordance with a previous study showing weak but specific immunoreactivity to anti-Olfm4 antibody in the Olfm4-L subset [2].

Out of the proteins that were significantly more abundant in the Olfm4-H or Olfm4-L neutrophils from healthy blood donors, 27 proteins other than Olfm4 displayed a bimodal distribution. Unfortunately, no surface protein with the potential to be used as a surrogate marker for one of the subsets was identified among these. Further, most of the bimodal proteins were detected at low levels and not in all the analysed blood donors. Several of them were also unrelated to the immune system. Reactome pathway enrichment analysis disclosed that both subsets showed significant enrichment in the neutrophil degranulation pathway, meaning that distinct proteins related to neutrophil granules are present in each subset. In the Olfm4-H cells, Rab3D, possibly involved in neutrophil degranulation [21], leucine rich alpha-2 glycoprotein, secreted from peroxidase-negative neutrophil granules and regulating myelopoiesis [22], and the V-type proton ATPase 16 kDa subunit, involved in acidification of neutrophil granules [23], showed higher abundance and were part of the neutrophil degranulation Reactome pathway. Meanwhile, the chemotactic alarmin S100-A7 [24], Rab3A, possibly involved in neutrophil degranulation [25], CXCR1, a major neutrophil chemotactic receptor that binds interleukin-8 [26], and neutrophil defensin alpha 4, an antimicrobial peptide with bactericidal activity [27], showed higher abundance in the Olfm4-L cells. Even though these proteins did not significantly colocalize to any subordinate Reactome pathway, the proteomic profiles of the subsets were distinct and involved proteins important for immunological and inflammatory processes in neutrophils. Interestingly, the Olfm4-H subset also showed higher abundance of CYBC1, a protein reported to be required for optimal production of the main subunit (gp91^*phox*^) of the neutrophil NAPDH oxidase and thus for the efficient production of reactive oxygen species (ROS) [28]. Meanwhile, the gp91^*phox*^ abundance did not differ between the subsets. Collectively, the data may suggest different preparedness of the subsets to respond to microbial stimuli. We [3] and others [29] have previously shown that in healthy individuals, the Olfm4-defined neutrophil subsets show equal propensity to phagocytose bacteria. In addition, the subsets display equal capacity to migrate to sterilely inflamed tissue and undergo apoptosis [3], but the present data indicate that more subtle differences in the mentioned processes may be present and these warrant attention in future studies.

To begin to investigate the role of the Olfm4-H neutrophils in septic shock, we initially planned to corroborate the correlation between the percentage of Olfm4-H neutrophils and disease severity in an adult population of septic shock patients [4, 6]. However, there was no significant correlation observed between severity of illness as measured by SOFA score or OSFD and the proportion of Olfm4-H neutrophils although there was such a trend regarding SOFA score. Further, the median percentage of Olfm4-H neutrophils reported in sepsis patients here was 20.5%, while the means in previous studies were approximately 25% in paediatric sepsis [4] and 26.6% in adult sepsis patients [6]. Using the published cut-off score of > 37.6% for independently predicting mortality in adult sepsis [6] only three of the 20 patients herein qualified. The sample size of the present study was based on previously published data on paediatric septic shock comparing patients with a complicated course of sepsis with a non-complicated course [4]. However, more recently published data showed a higher variation in the percentage of Olfm4-H neutrophils in adult septic shock patients as well as a smaller difference in percentage between patients who survived and died [6]. Thus, the present study was underpowered for corroborating the published findings, probably explaining the lack of association between clinical findings and the findings from proteomics discussed below. On the other hand, the data confirmed that the plasma concentration of Olfm4 was significantly higher in sepsis patients than in healthy controls, as previously reported for children [4], and the increased concentration correlated with neutrophil concentration in the blood.

Having discovered different proteomic profiles in the Olfm4-defined subsets at baseline and based on previous larger studies showing that a high percentage of Olfm4-H neutrophil predicts a worse outcome in sepsis, we hypothesized that it is the distinct profile of the Olfm4-H neutrophils that predispose the cells to detrimental responses during infection that may contribute to sepsis pathogenesis. As expected, using LC-MS/MS analysis of FACS-sorted Olfm4-H and Olfm4-L neutrophils from septic shock patients, the data quality and most abundant proteins found corresponded to those from neutrophils isolated from healthy blood donors. Again, Olfm4 was the only major granule protein with a significantly increased abundance in one subset. However, the subsets again displayed distinct proteomic profiles, encompassing partly different proteins as compared to neutrophils from healthy donors. Interestingly, the Olfm4-H cells in the septic shock patients, like in the healthy blood donors, showed increased abundance of CYBC1, important for ROS production [28], but also of NLR family member X1, involved in the regulation of several components of immune system function [30]. On the other hand, the Olfm4-L subset displayed increased abundance of Fc epsilon receptor 1 gamma subunit, involved in immunoreceptor signal transduction [31], the low-affinity IgG receptor Fc gamma receptor IIa which activates neutrophils upon cross-linking [32], WASH complex subunit 4, important for actin reorganization in endosomal trafficking [33], and the MHC class I component HLA class I histocompatibility antigen A alpha chain, essential for endogenous and viral antigen presentation to CD8+ T cells [34]. A previous study identified decreased surface expression of CD64 on Olfm4-H relative to Olfm4-L neutrophils in children after bone morrow transplantation [7], a finding that was not reflected in the whole-cell material analysed here.

Reactome pathway analysis showed near-significant enrichment in the Olfm4-L subset in the pathways neutrophil degranulation and antigen processing – cross presentation, the latter related to presentation through MHC class I. Meanwhile, there was no enrichment in the neutrophil degranulation pathway in the Olfm4-H subset. Previous studies regarding neutrophil Olfm4 functions during infection have been carried out in mice. It was discovered more than ten years ago that mice lacking Olfm4 show reduced colonization with *Helicobacter pylori* and enhanced activation of the nuclear factor-kappaB activity induced by the bacteria as compared to wild-type mice [35]. Later it was demonstrated that Olfm4 inhibits cathepsin C-mediated protease activities, and Olfm4-decifient mice showed enhanced capacity to kill *Staphylococcus aureus* and *Eschirichia coli* as compared to wild-type mice [36]. The same group finally showed that Olfm4-deficient mouse neutrophils display reduced hydrogen peroxide-induced apoptosis as compared to neutrophils from wild-type mice [37]. In 2018 it was discovered that mouse neutrophils also display a bimodal distribution of Olfm4 [5]. Olfm4-expressing mice displayed more severe mucosal damage than Olfm4-deficient mice in a model of intestinal ischemia-reperfusion injury, with increased nuclear factor-kappaB and inducible nitric oxygen synthase activation, in a neutrophil-mediated manner [38]. Recently, it was shown that juvenile Olfm4-decifient mice are protected from sepsis, and display lower levels of acute kidney injury than in wild-type mice [18]. All the mentioned studies are consistent with the idea of detrimental processes in Olfm4-containing neutrophils during infection, with defective bacterial clearance or an exaggerated pro-inflammatory response. On the other hand, Kangelaris *et al.* recently published data suggesting that the Olfm4-H neutrophil percentage predicts death also in non-sepsis systemic inflammatory respiratory syndrome (SIRS), which in combination with data showing no association between an elevated Olfm4-H percentage and culture positivity or other biomarkers of poor sepsis prognosis, led the authors to suggest a non-microbial mechanism being responsible for the ability of Olfm4-percentage to predict death in critically ill patients [6]. The data presented herein support different responses of the neutrophil subsets during sepsis in several pathways related to infection and inflammation, and further studies are needed to delineate which of the proteins and pathways are pivotal for the detrimental outcome related to Olfm4-H neutrophils.

In conclusion, the human Olfm4-defined neutrophil subsets display distinct proteomic profiles both at baseline, with Olfm4-L neutrophils showing increased abundance in several proteins that may enhance the response to infection, and in septic shock where the Olfm4-L subset displays increased abundance of proteins related to activation by immunoglobulins. This is the first report pointing towards specific differential functions of the Olfm4-defined neutrophil subpopulations, and the results are in line with previous findings of a distinct role of Olfm4-H neutrophils in bacterial infection and under proinflammatory conditions. The findings incite further investigations into the role of the revealed proteomic profiles in the individual subsets in the pathogenesis of sepsis.

## Statements

## Acknowledgements

We are grateful to study nurses Helén Didriksson and Carina Jonsson for arranging the patient samples. Flow cytometry was performed using instruments at the Flow Cytometry Core Facility, Faculty of Medicine and Healthy Sciences, Linköping University. We would like to acknowledge the Proteomics Core Facility, Faculty of Medicine and Health Sciences, Linköping University for assistance with proteomics and the Bioinformatics Core Facility, Faculty of Medicine and Health Sciences and Clinical Genomics Linköping, Science for Life Laboratory, Department of Biomedical and Clinical Sciences, Linköping University for assistance with bioinformatics analyses.

## Statement of ethics

Peripheral blood from healthy blood donors was obtained through the Linköping University Hospital blood bank. The donors had given written consent that blood could be used for research purposes and only de-identified samples were received. In accordance with the Declaration of Helsinki and paragraph 4 of the Swedish law (2003:460) on Ethical Conduct in Human Research, no specific ethical approval is needed. Blood samples from septic shock patients was obtained after approval of the study protocol by the regional ethical review board (Linköping 2016/361-31, 2018/631-32) and with informed written consent from the patient or next of kin.

## Author contributions

A.W. conceived the study, planned the experiments, analysed the data, and wrote the manuscript together with H.L and with input from all authors. In addition, H.L. performed experiments and analysed the data. H.A. and M.C. provided access to patient samples and clinical data and contributed to the study design and data analysis. J.D. performed and contributed to interpretation of bioinformatic analysis. M.T. contributed to development of the proteomics protocol, performed LC-MS/MS and analysed proteomics data. All authors read and approved the final manuscript.

## Data availability statement

All data generated or analyzed during this study are included in this article and its supplementary material files. Further enquiries can be directed to the corresponding author.

