## Supplementary figures and tables for "The Olfm4-defined human neutrophil subsets differ in proteomic profile in septic shock": Supplementary material.pdf

**Supplementary figure legends**

**Supplementary figure S1.** *Gating strategy for fluorescence-activated cell sorting (FACS) of Olfm4-defined neutrophil subsets.* Granulocytes were isolated, fixed, permeabilized and stained with antibodies as described in the main text. Neutrophils were identified based on high CD15-staining and sorted into the two populations Olfm4-H and Olfm4-L.

**Supplementary figure S2.** *Effect of whole blood incubation on the proportion of Olfm4-H neutrophils and plasma Olfm4 concentration.* Healthy blood donor whole blood samples were either incubated at room temperature for 20 hours or processed immediately. Olfm4 staining and plasma Olfm4 measurements were performed as described in the main text. Effect of delayed processing on the proportion of Olfm4-H neutrophils (A) and plasma Olfm4 concentrations (B) are shown. Statistical analysis of differences between groups was performed by Wilcoxon signed rank test (paired data), showing no significant differences (n=3).

**Supplementary figure S3.** *Imaging flow cytometry gating strategy to quantify the proportion of Olfm4-H neutrophils.* Whole blood leukocytes were fixed, permeabilized and stained with anti-CD15 and anti-Olfm4 antibodies as well as DAPI and analyzed by imaging flow cytometry (ImageStreamX MK II) as described in the main text. The cells were gated in 5 steps (A-E) sequentially. Examples of gated and non-gated cells are shown with arrows. (A) The cells in focus were gated based on gradient RMS in the brightfield channel. (B) Single cells were gated based on area versus aspect ratio in the brightfield channel. (C) Cells positive for the nuclear stain DAPI (excited by the 405 nm laser) were gated. (D) Neutrophils were gated based on

CD15-PE staining (excited by the 488 nm laser) and fluorescence intensity in the side scatter channel (autofluorescence upon excitation with the 785 nm laser). (E) Finally, high fluorescence intensity in the Alexa Fluor 647 channel (excited by the 642 nm laser) identified Olfm4-H neutrophils while low intensity in the same channel identified the OLFM4-L neutrophils. The gate for OLFM4-H cells was validated for each individual donor using a control sample where the anti-OLFM4-serum was omitted.

**Supplementary tables**

**Supplementary table S1.** Proteins identified by LC-MS/MS in the Olfm4-defined neutrophil subsets from healthy blood donors.

**Supplementary table S2.** Reactome pathway analysis of differentially abundant proteins in the Olfm4-defined subsets in healthy blood donors.

**Supplementary table S3.** Proteins identified by LC-MS/MS in the Olfm4-defined neutrophil subsets from septic shock patients.

**Supplementary table S4.** Reactome pathway analysis of differentially abundant proteins in the Olfm4-defined subsets in septic shock patients.
