## Supplementary figures and images for "The Olfm4-defined human neutrophil subsets differ in proteomic profile in septic shock"

### Fig S1.pdf

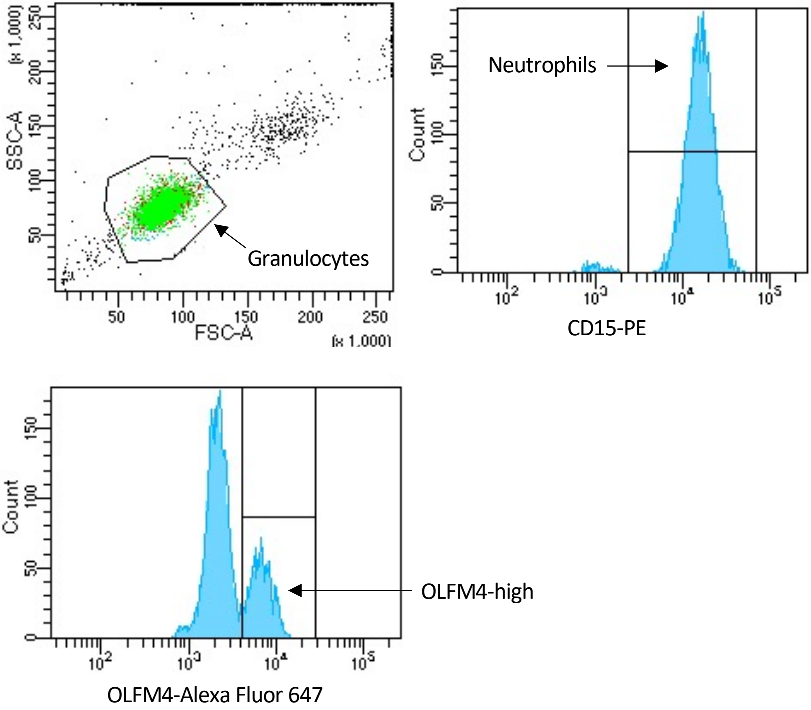

### Fig S2.pdf

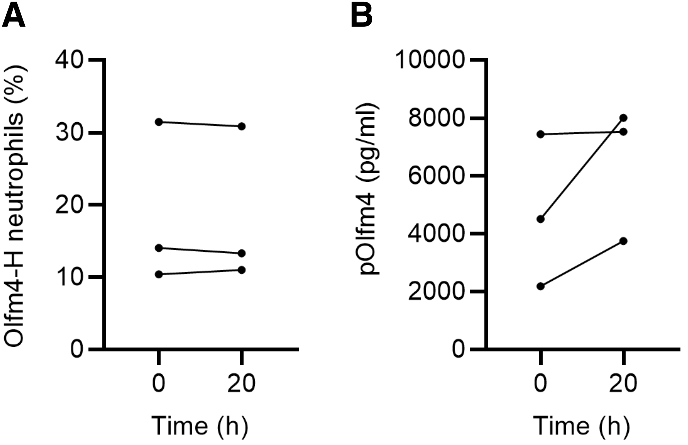

### Fig S3.pdf

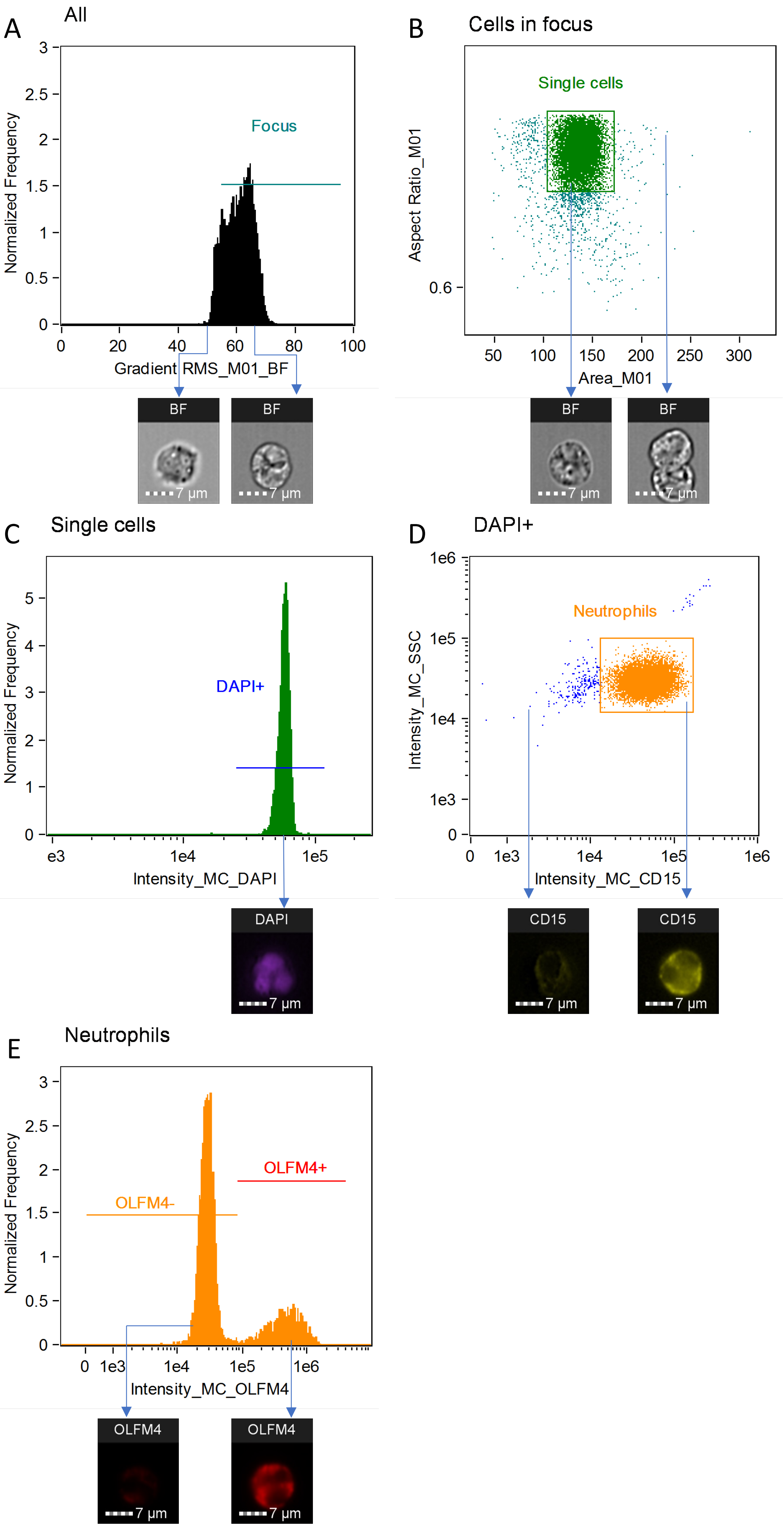
